# Fibroblast Growth Factor 5 (FGF5) and Its Missense Mutant FGF5-H174 Underlying Trichomegaly: A Molecular Dynamics Simulation Investigation

**DOI:** 10.1101/2022.12.10.519912

**Authors:** Skyler H. Hoang

## Abstract

The missense mutation Y174H of FGF5 (FGF5-H174) had been associated with trichomegaly, characterized by abnormally long and pigmented eyelashes. The amino acid tyrosine (Tyr/Y) is conserved across many species, proposedly holding important characteristics for the functions of FGF5. One-microsecond molecular dynamics simulations along with protein-protein docking and residue interacting network analysis were employed to investigate the structural dynamics and binding mode of both wild-type (FGF5-WT) and its mutated counterpart (FGF5-Y174H). It was found that the mutation caused decreases in number of hydrogen bonds within the protein, sheet secondary structure, interaction of residues 174 with others, and number of salt-bridges. On the other hand, the mutation showed increases in solvent accessible surface area, number of hydrogen bonds between the protein and solvent, coil secondary structure, protein C-alpha backbone root mean square deviation, protein residue root mean square fluctuations, as well as occupied conformational space. In addition, protein-protein docking integrated with molecular dynamics simulations and molecular mechanics - Poisson-Boltzmann surface area (MM/PBSA) binding energy calculation demonstrated that the mutated variant possessed stronger binding affinity towards fibroblast growth factor receptor 1 (FGFR1). However, residue interaction network analysis demonstrated that the binding mode was drastically different from that of the FGF5-WT-FGFR1 complex. In conclusion, the missense mutation conferred more instability, stronger binding affinity towards FGFR1 but with distinctively altered binding mode or residue connectivity. These findings might help explain the decreased activation of FGFR1, underlying trichomegaly.

## Introduction

Fibroblast growth factor (FGF) 5 (FGF5), belonging to the canonical FGF family as well as FGF4 subfamily, has a substantial impact on the regulation of cell differentiation and proliferation (Ornitz & Itoh, 2015). Particularly, FGF5 is most notable for its function in the regulation hair growth: by activating the fibroblast growth factor receptor 1 (FGFR1) and being predominantly produced in hair follicles, it acts as an inhibitor of hair elongation by promoting the transition from the hair follicle’s growth phase, anagen, into the apoptosis-induced regression phase, catagen (Higgins et al., 2014). Simply put, it is required for the proper regulation of the hair development cycle. In terms of direct application, CRISPR/Cas9-mediated loss of FGF5 function promotes the production of longer wool in Chinese merino sheep, representing a valuable target for the wool industry (Li et al., 2017).

Other *in vitro*, *in vivo*, and clinical studies have demonstrated other functions of FGF5. In terms of molecular signaling, the FGF-FGFR signaling motif frequently depends on the phosphoinositide-3-kinase–protein kinase B/Akt (PI3K-AKT), rat sarcoma mitogen-activated protein kinase (RAS-MAPK), and signal transducer and activator of transcription (STAT) pathways, master regulators of many biological processes (Ornitz & Itoh, 2015). Particularly, in an *in vitro* cultured primary rat Schwann cells, FGF5 was demonstrated to be an autocrine regulator of Schwann cells that could regulate Schwann cell migration and adhesion through inhibiting extracellular signal-regulated protein kinases 1 and 2 (ERK1/2) MAPK activity and the upregulation of N-cadherin (B. Chen et al., 2020). Additionally, in a study with *in vitro* and clinical combined approach, FGF5 was found to be overexpressed in osteosarcoma cell lines and clinical tissue samples and to promote osteosarcoma cell proliferation by activating the MAPK signaling pathway, suggesting that FGF5 could be a therapeutic target for osteosarcoma (Han et al., 2019). On the other hand, in a study with *in vitro* and *in vivo* combined approach, FGF5 can contribute to the malignant progression of human astrocytic brain tumors by stimulating proliferation, migration, and tube formation in human umbilical vein endothelial cells via autocrine and paracrine effects (Allerstorfer et al., 2008). Additionally, an *in vitro* study showed that methylated exogenous FGF5 overexpression conferred resistance to cisplatin treatment; meanwhile, FGF5 was not expressed in non-cancerous esophageal tissues, and only a small percentage of cells had it methylated, indicating that FGF5 methylation could be a sensitivity marker of esophageal squamous cell carcinoma to definitive chemoradiotherapy a biomarker (Iwabu et al., 2019). From a clinical perspective, a recent clinical study demonstrated that FGF5 may regulate to blood pressure, as patients with primary hypertension had abnormally high levels of FGF5 mRNA and protein expression, along with genetic variations that were related to their blood pressure (Ren et al., 2018). In conjunction, another clinical found concluded that the genetic variant rs148038 near FGF5 gene, highly present in the Filipino population, was strongly associated with blood pressure response to calcium channels blockers (Punzalan et al., 2022). A case-control association analysis found that the FGF5 polymorphism rs16998073 significantly increases the risk of preeclampsia in the Chinese Han population, suggesting that FGF5 may play a role in the progression of the disease (Xin et al., 2022). Overall, emerging explorations into the functions of FGF5 emphasizes the need for further research into this unique protein.

Recent evidence suggested that Y174 is a highly conserved residue across species possessing critical roles in structure formation and functions of FGF5. The missense mutation of this residue into histidine (Y174H) may the underlying cause of trichomegaly, a disorder characterized by excessively long, pigmented, and thick eyelashes (Higgins et al., 2014; Rossetto et al., 2013). Y174 is characterized by its hydrophobic aromatic function group, whereas H174 is characterized by a positively charged imidazole functional group (**Figure 1A & B, top panels**). This poses a question: will such a drastic difference interfere with structures and functions of FGF5? Herein, *in silico* approaches (i.e., molecular dynamics simulations and protein-protein docking) were used to test the hypothesis that the missense mutation could lead to severe conformational changes and protein instability, resulting in reduced binding at FGFR1 and the development of trichomegaly.

**Figure 1.**
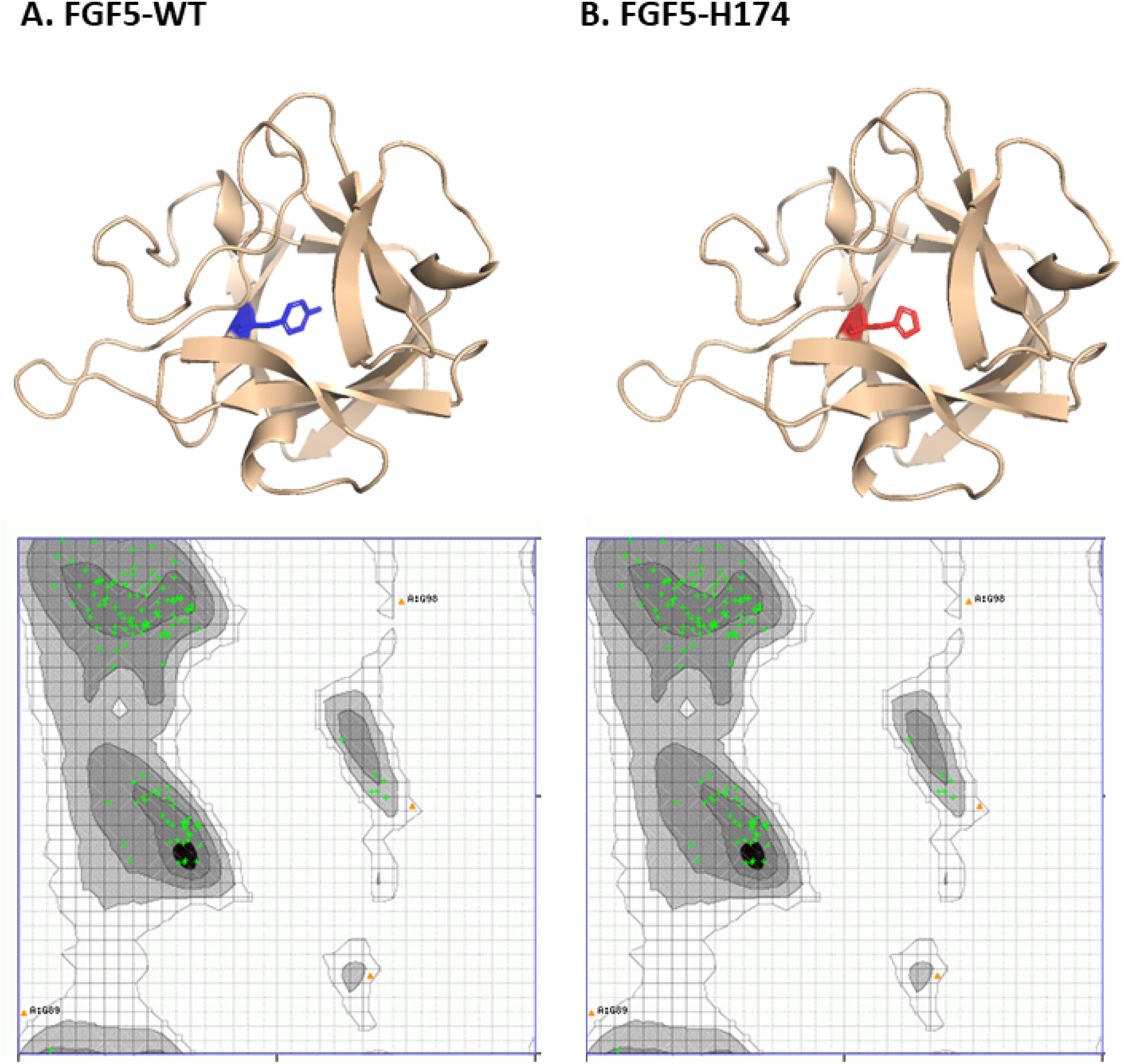
Structures of (A) FGF5-WT and (B) FGF5-H174 with their respective Ramachandran plots.

## Method

### 1. Protein Retrieval and *in silico* mutagenesis

The putative structural model of FGF5 was retrieved from AlphaFold Protein Structure

Database (https://alphafold.ebi.ac.uk/), with accession number: P12034. PyMol 2.5 (https://pymol.org/2/) was used to perform the mutagenesis at 174 from tyrosine (Y174) to histidine (H174); the conformation with the least steric strain was applied. PyMol 2.5 was also used to truncate two residue segments, which were N-terminus to residue 83, and residue 220 to C-terminus. These segments contained residues with low or very low confidence scores, which could be confounding factors to structural and dynamics investigations using molecular dynamics simulations. Ramachandran Plot Server (https://zlab.umassmed.edu/bu/rama/) was employed to validate the models; both showed 139 (97.015%) highly preferred and 4 (2.985%) preferred observations with no questionable observation (**Figure 1A & B, bottom panels**).

### 2. Molecular Dynamics Simulation setup

YASARA Dynamics (Version 21.8.27) (Land & Humble, 2018) was used to construct 2000-nanosecond molecular dynamics simulations with 2,001 simulation snapshots each for FGF5 proteins and 200-nanosecond molecular dynamics simulations with 1,001 simulation snapshots for FGFR1-FGF5 complexes. The script md_run.mcr was the backbone of the simulations. In brief, simulation were optimized with: (1) steepest descent minimization without electrostatics (2) simulated annealing minimization of the solvent (3) simulated annealing of the solvent to adapt to deleted waters (4) running 50 steps of solvent MD (5) running local steepest descent minimization without electrostatics to remove bumps. Timestep settings were set at the followings: 1.25 femtosecond of intramolecular force substep and intermolecular forces are calculated for every 2.5 femtosecond. Concentration of NaCl was set at 0.9%; pH was set at 7.4. Pressure control was in Solvent Probe mode. Extension of the simulation cell is 20 Å larger than the protein. Shape of the simulation cell is cube. The forcefield of the simulations was adapted AMBER14 with OL15 DNA fine-tuning with TIP3P water model (Galindo-Murillo et al., 2016). Simulations were performed with long-range Coulomb forces (particle-mesh Ewald). “CorrectDrift” mode was turned on to keep solute from diffusing around and crossing the periodic boundaries. Afterwards, md_analyze.mcr script was used to retrieve data for (1) solvent accessible surface area (SASA), (2) percentages of types of secondary structures throughout the simulations, (3) interactions between residue 174 and others, (4) root mean square deviation (RMSD) of C-alpha carbons of the proteins, (5) number of salt-bridges created by Lysine (Lys)/Arginine (Arg) and Aspartic acid (Asp)/Glutamic acid (Glu), protein residue root mean square fluctuations (RMSF). The md_analyebindenergy.mcr script was used to retrieve molecular mechanics – Poisson-Boltzmann-derived binding energy (MM/PBSA). It was calculated based on the following premise:

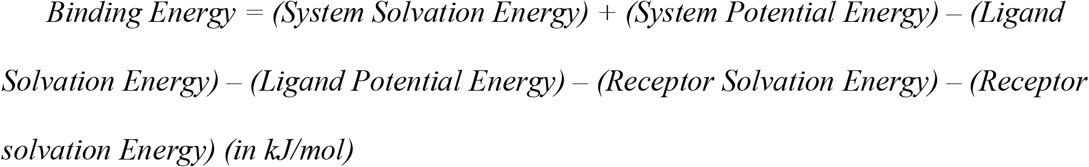

In which, more negative binding energies indicate better binding; more positive energies do not indicate no binding.

### 3. Construction of PCA free energy landscapes

The YASARA molecular dynamics simulation trajectory.sim files were then converted to GROMACS simulation.xtc files using md_convert.mcr script. The gmx covar, gmx anaeig, gmx sham functions in GROMACS 2022.4 were then used to retrieve principal component analysis (PCA) of the movement of C-alpha backbone of the proteins, which was eventually used to construct free energy landscapes and extract meta-stable states.

### 4. Protein-protein docking and constructing residue interacting network

Protein-protein docking was performed with HDock Server (Yan et al., 2017) for FGF5 proteins in their meta-stable states towards the FGFR1 receptor (PDB 1FQ9:C). The models with the highest scoring functions were used in this study. The protein-protein complexes were visualized and structurally analyzed with PyMol, and residue interactions analysis were created in the formed of network through Residue Interaction Network Server 3.0 (Clementel et al., 2022). Cytoscape 3.9.1 (https://cytoscape.org) was used to calculate degree, which counts for the number of edges or interactions a residue can form. GraphPad Prism 9 ((https://www.graphpad.com/), was used to create histograms for the degree.

### 5. Visualization

Data visualization was accomplished with PyMol, GraphPad Prism 9, YASARA, QtGrace (https://sourceforge.net/projects/qtgrace/), and Cytoscape.

## Results

### 1. The missense mutation Y174H altered interactions with the solvent

Parameters relevant to protein-solvent interactions (solvent accessible surface are, number of hydrogen bonds within the protein, and between protein and solvent) were evaluated (**Figure 2**). Particularly, on average, FGF5-H174 with 7150 ± 139.3 Å^2^ showed higher solvent accessible surface area than FGF5-WT with 7342 ± 137.3 Å^2^ (**Figure 2A**). FGF5-WT demonstrated apparent increase in solvent accessible surface area from approximately 200^th^ ns to 300^th^ ns and 700^th^ ns to 1200^th^ ns (**Figure 2A**). In addition, the number of hydrogen bonds within FGF5-H174 was 95.3 ± 5.13, which was smaller than that of FGF5-WT (97.94 ± 4.95 hydrogen bonds) (**Figure 2B**). Specifically, the difference was most apparent at the 200^th^ ns to the 300^th^ ns and at 1100^th^ ns to 1600^th^ ns (**Figure 2B**). Alternatively, in conjunction with the decrease in intramolecular hydrogen bonds, the number of hydrogen bonds of FGF5-H174 with its solvent was 256.3 ± 9.45, which was larger than that of FGF5-WT with 249.5 ± 8.99 hydrogen bonds (**Figure 2C**). The difference was also most apparent at the 200^th^ ns to the 300^th^ ns and at 1100^th^ ns to 1600^th^ ns (**Figure 2C**).

**Figure 2.**
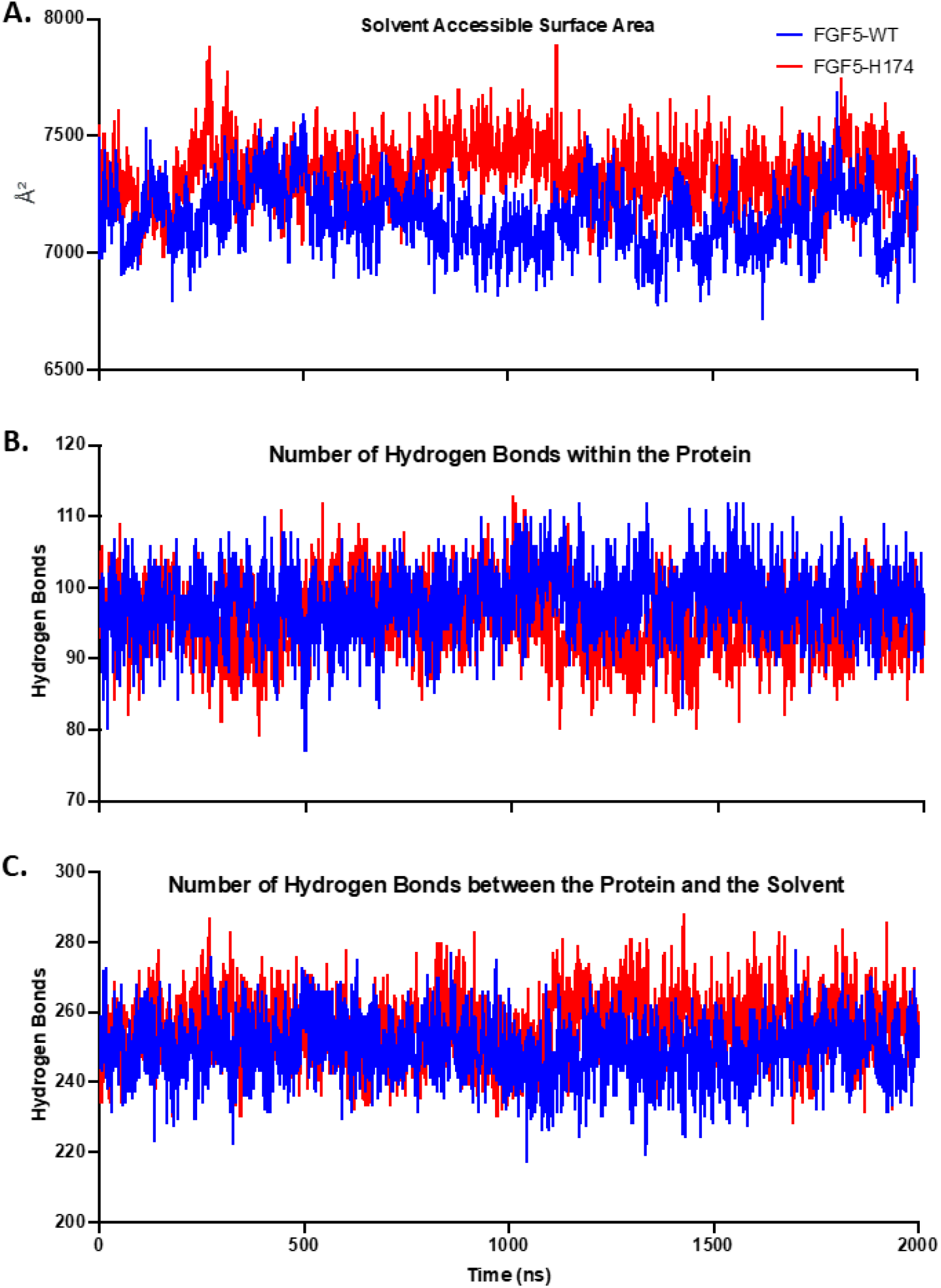
Behavioral parameters of wild-type fibroblast growth factor 5 (FGF5-WT) and its Y174H missense mutation variant (FGF5-H174). (A) Solvent accessible surface areas (SASA) in Å^2^. (B) Number of hydrogen bonds within the proteins. (C) Number of hydrogen bonds between the protein and solvent

### 2. The missense mutation Y174H decreased sheet and increases coil secondary structures

Parameters relevant to the secondary structures of (percentages of sheet, turn, and coil structures) were evaluated (**Figure 3**). Particularly, due to the mutation, the percentage of sheet structure was slightly decreased from 51.57% ± 3.622% to 49.76% ± 3.121% for FGF5-WT and FGF5-Y174H, respectively (**Figure 3A**); turn structure were similar with 24.56% ± 2.086% and 23.84% ± 2.413% for FGF5-WT and FGF5-H174, respectively (**Figure 3B**), and coil structure was slightly increased from 23.87% ± 3.730% to 26.39% ± 3.585% (for FGF5-WT and FGF5-Y174H, respectively) (**Figure 3C**). Secondary structures were also presented as time-series graphs (**Figure 3D** and **3E**). From the 260^th^ ns and beyond, turn and sheet structures at residues 146 to 152 (**Figure 3D**) transformed into coil structure due to the mutation (**Figure 3E**), or from the 170^th^ ns and beyond, between residues 160 to 171, sheet structure was decreased and substituted with coil structure.

**Figure 3.**
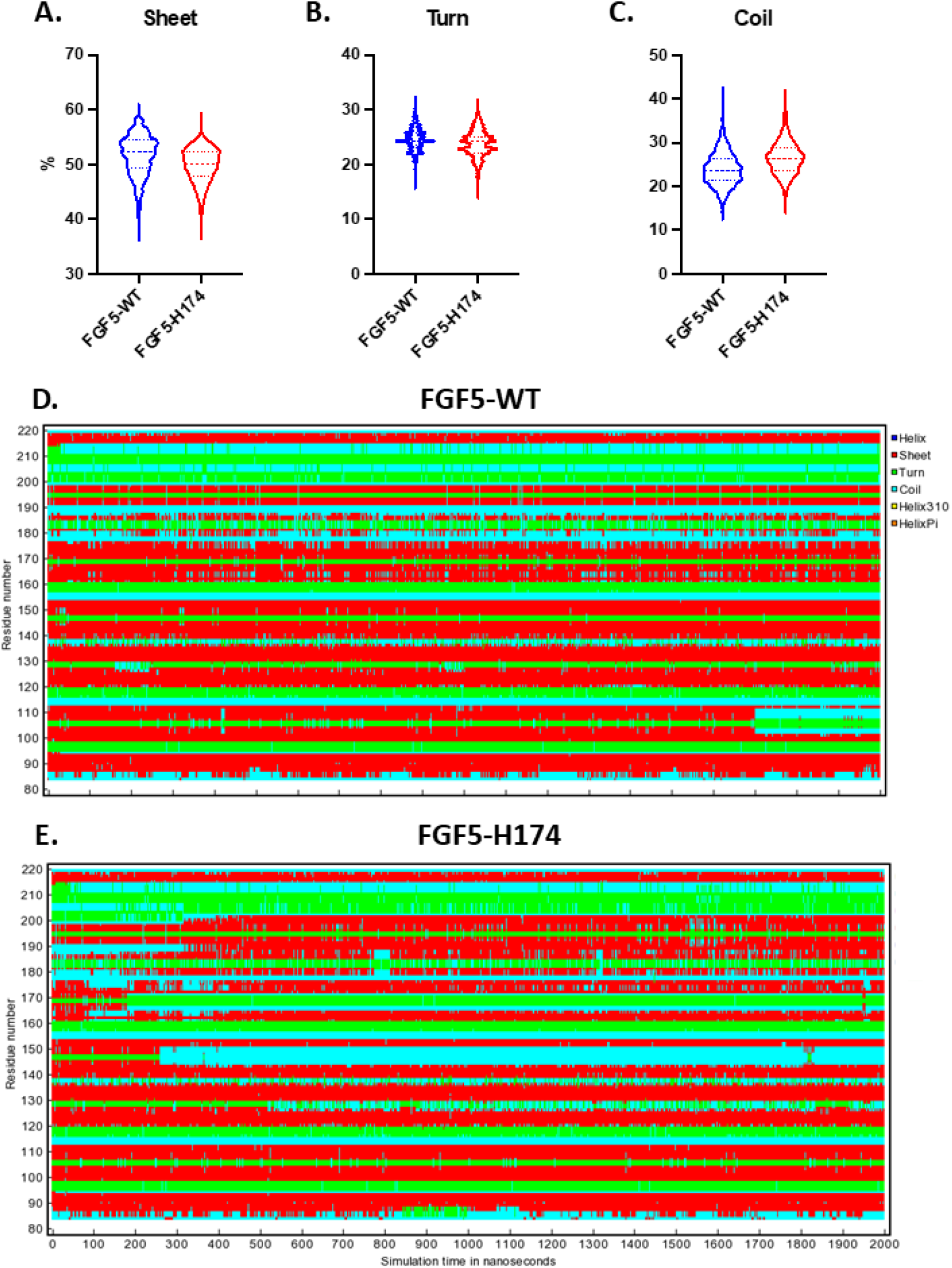
Assessment of secondary structure assessment over time. Violin graphs depicting the distributions of (A) sheet (B) turn (C) coil secondary structures over time. Color-filled time-series graphs depicting the evolutions of these structures over time for (D) FGF5-WT and (E) FGF5-H174.

### 3. Decreased interaction of residue 174 with other residues was observed for H174

Interactions of both wild type (Y174) and mutated (H174) residues at position 174 with other residues were evaluated (**Figure 4**). Both were able to form interactions with the following residues: 91, 123, 131, 133, 162, 164, 172, 190, and 215, with the exception that the mutated H174 residue could form a short-lived hydrogen bond with residue 163 (**Figure 4A** and **4B**). Overall, there was a decrease in interacting time for all the common residues for H174 and the hydrogen bond between Y174H and 163 was short-lived, as it mostly lasted only between the initiation of the stimulation up to approximately the 370^th^ ns of the simulation (**Figure 4**). In addition, most interactions with other residues for Y174 were sustained throughout the simulations, whereas residues 131, 133, 162, 164, and 190 were sporadically sustained (**Figure 4**).

**Figure 4.**
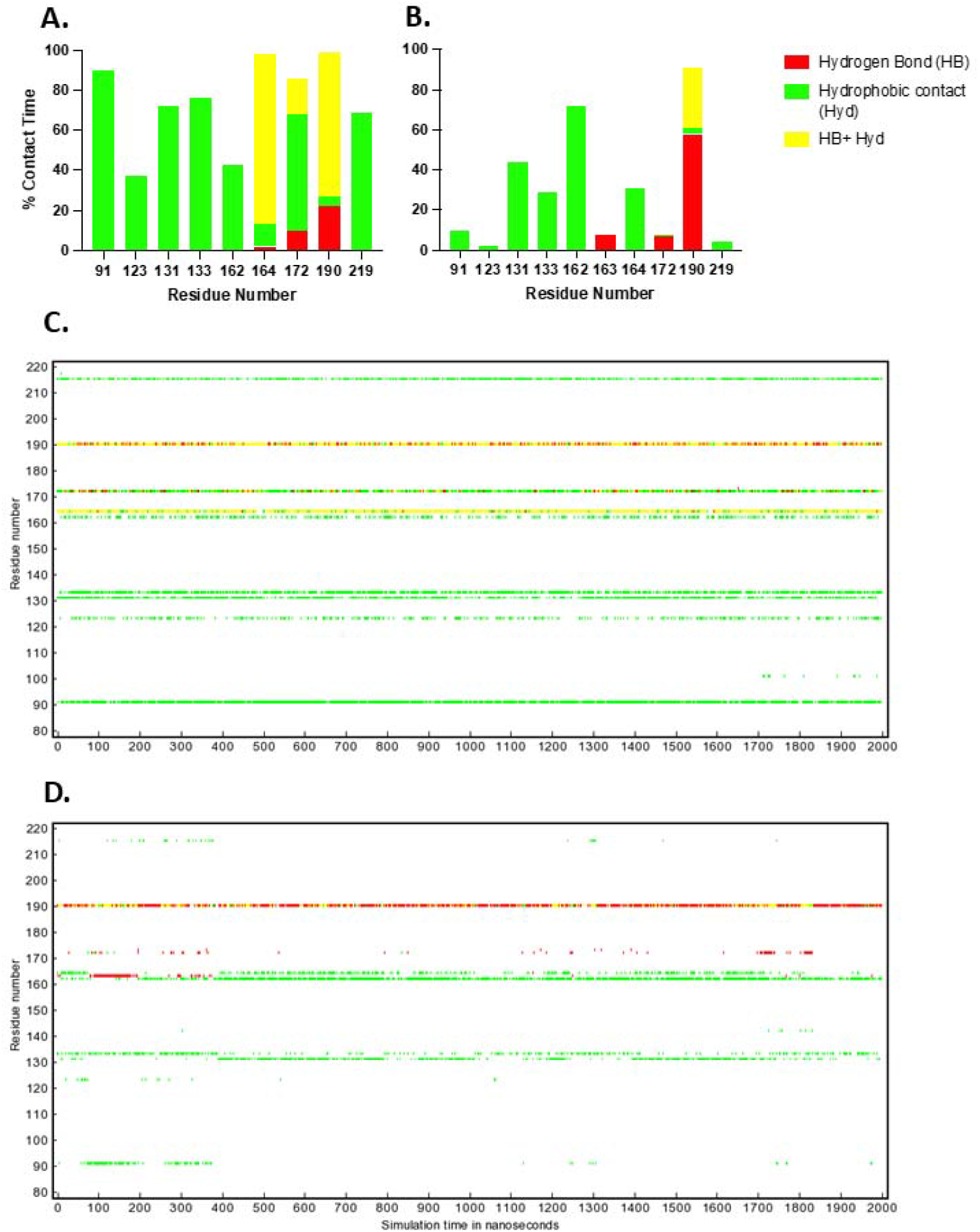
Analyses of interactions of Y174 and H174 with other residues. Bar graphs depicting the accumulations of interactions of residue 174 with others for (A) the wild-type Y174 and (B) the missense mutation variant H174. Color-filled time-series graphs depicting the evolutions of these interactions over time for (C) Y174 and (D) H174; white space denotes no interaction.

### 4. FGF5-Y174H experienced more movement, instability, conformational changes

To compare the movement, stability, and conformations between the wild-type and mutated proteins, data regarding C-alpha backbone RMSD, number of salt-bridges, residue RMSF (**Figure 5A)** were evaluated. FGF5-H174 with 1.771 ± 0.3027 Å demonstrated more conformational changes as well as movement throughout the simulations than FGF5-WT with 1.195 ± 0.1706 Å (**Figure 5A**). Nevertheless, both RMSD values were smaller than 2 Å, which indicated that the starting structures were well predicted and validly constructed. The C-alpha RMSD of FGF5-WT was mostly converging with only a small spike at around the 500^th^ of the simulation, whereas FGF5-Y174H experienced a gradual increase in RMSD up until the approximately the 1100^th^ ns and did not reach convergence until the 1200^th^ ns (**Figure 5A**). Inversely correlated with the trends of C-alpha RMSDs, FGF5-WT had 6.153 ± 0.9714 salt bridges, higher than FGF5-Y174H with 5.228 ± 1.040 salt bridges, and an apparent decrease in the number of salt-bridges could also be observed at around the 500^th^ ns (**Figure 5B**). Additionally, the RMSF for FGF5-Y174H with 1.387 ± 0.8201 Å was higher than that of FGF5-WT 1.171 ± 0.6581 Å (**Figure 5C**). Specifically, the 3 regions with the most apparent increases for FGF5-Y174H include: residues 143 to 152, 162 to 175, and 196 to 213 (**Figure 5C**).

**Figure 5.**
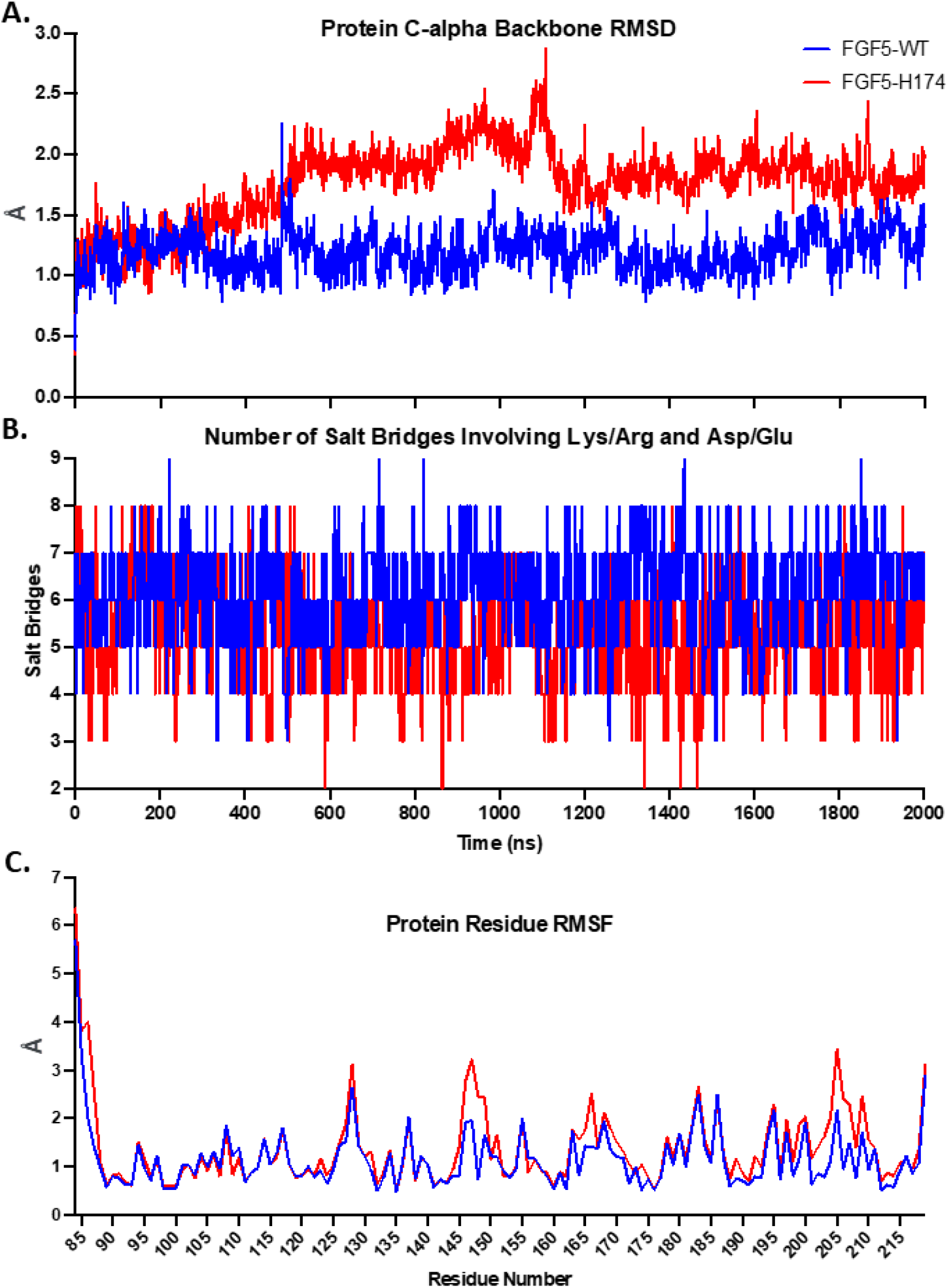
Stability and conformation parameters of FGF5-WT and FGF5-Y174H. (A) Root mean-square deviation (RMSD) of C-alpha backbones. (B) Number of salt-bridges formed by the interactions of lysine (Lys)/ arginine (Arg) with aspartic acid (Asp)/ glutamic acid (Glu). (C) Root mean-square fluctuations (RMSF).

These findings warranted further examination and validation with the free energy landscapes constructed by applying Boltzmann-inverting multi-dimensional histogram on the PCA of the movement of C-alpha backbone. The free energy landscapes demonstrated that the conformational movement of FGF5-H174H was more scattered and occupied more space than FGF5-WT (**Figure 6A and B, top panels**). In conjunction, it was predicted that it would cost 8.02 kJ/mol and 9.17 kJ/mol Gibbs free energies for FGF5-WT and FGF5-H174 (respectively) to reach meta-stable states from the high-energy states, supporting the idea that FGF5-H174 might experience more movement and conformational changes (**Figure 6A and B, top panels**). Meta-stables were also extracted for visualization (**Figure 6A and B, bottom panels**) and structural alignment analysis (**Figure 7**). Particularly, the missense mutation induced apparent confirmational shifts to 4 loop segments of the protein, which were: K146 to K149, S127 to I130, R200 to H210, as well as Q167 to Y171 (**Figure 7**). These findings supported the speculation that such drastic conformational drifts derived from the missense mutation might reduce pharmacological activity or binding affinity of FGF5 at FGFR1.

**Figure 6.**
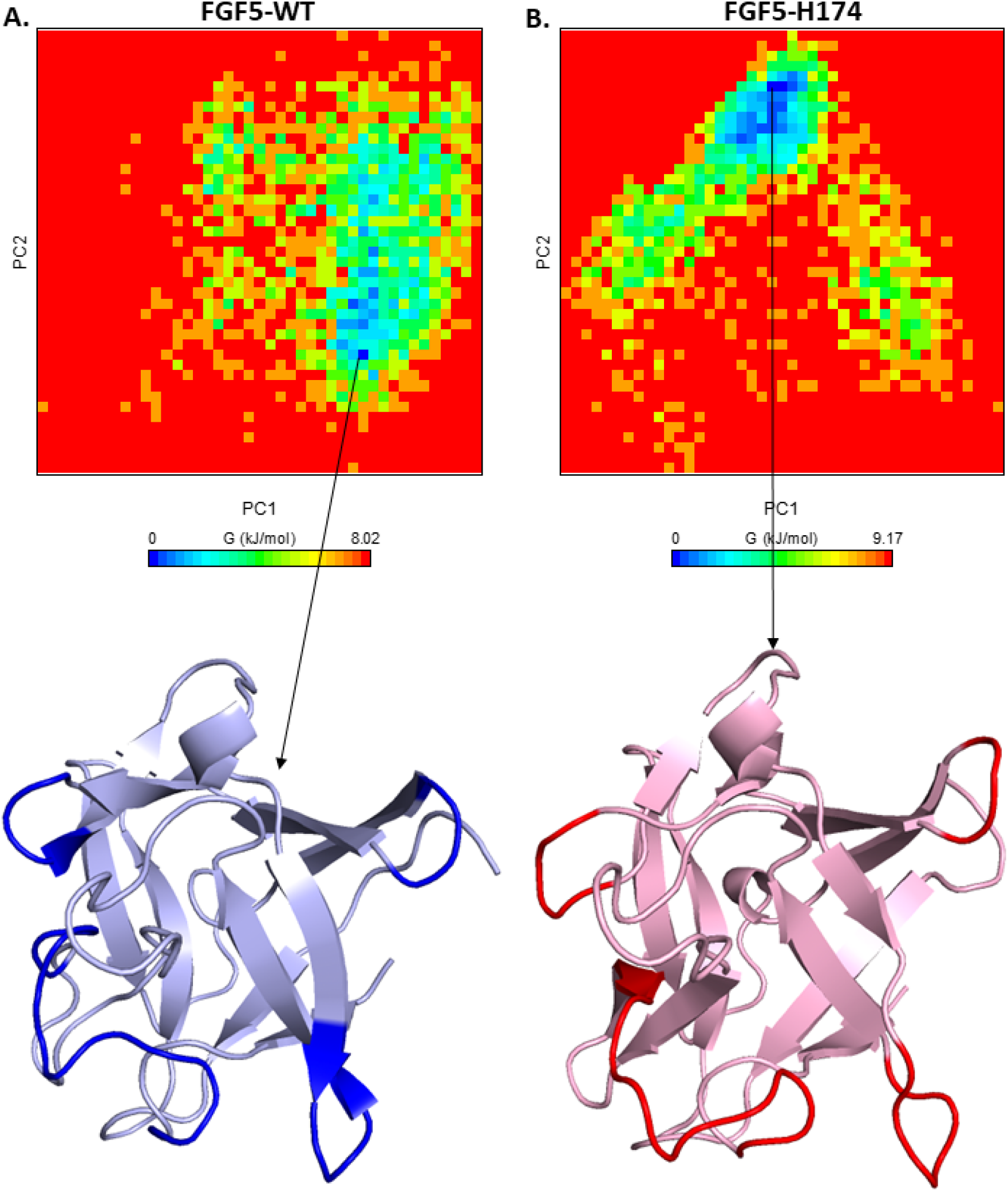
Free energy landscapes of FGF5-WT and FGF5-H174 with meta-stable states derived from the C-alpha backbone principal component analysis (PCA). (A) FGF5-WT (B) FGF5-H174. Bold colors denoted drastic conformational changes.

**Figure 7.**
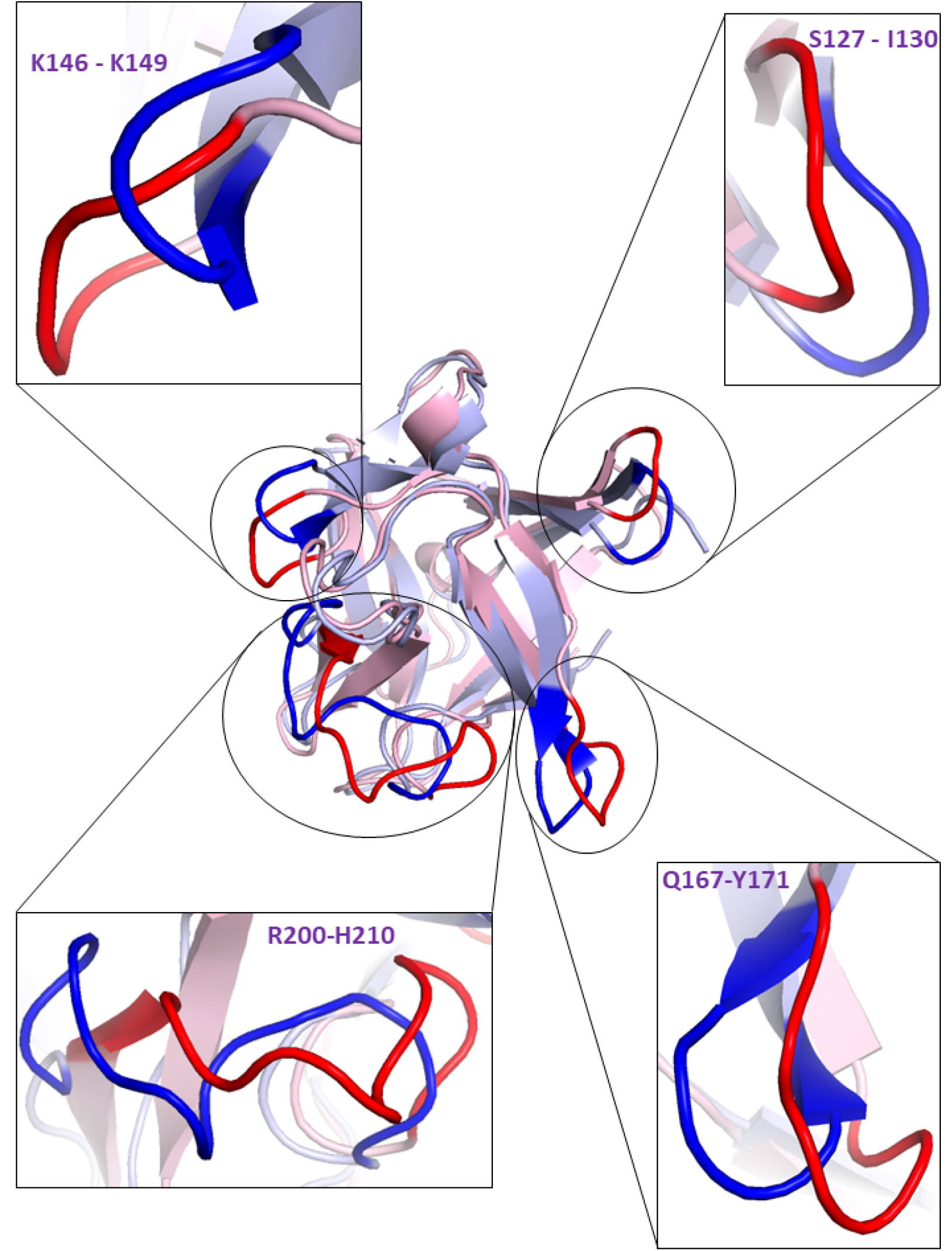
Structural alignment of meta-stable FGF5-WT (blue) and FGF5-H174 (red). Bold colors denoted drastic conformational changes.

### 5. The mutation increased binding affinities towards FGFR1 with altered binding mode

To examine such speculation, protein-protein docking was performed for FGF5 proteins in meta-stable states and FGFR1; molecular dynamics simulations with MM/PBSA binding energy calculations for the complexes were subsequently performed (**Figure 8**). Contrary to the hypothesis that the drastic conformational shifts derived from the missense mutation would reduce binding affinity, the FGFR1 - FGF5-H174 complex not only had lower C-alpha backbone RMSD (of 2.896 ± 0.4400 Å) that that of FGFR1 - FGF5-WT (3.440 ± 0.4636 Å), but also FGFR1 - FGF5-H174 complex reached convergence sooner (at around the 100^th^ ns) than FGFR1-FGF5-WT complex, which did not reach convergence until the 125^th^ ns (**Figure 8A**). The RMSDs for FGFR1 - FGF5-WT and - FGF5-H174 calculated from the 125^th^ ns were 3.44 ± 0.36 Å and 2.70 ± 0.30 Å, respectively (**Figure 8A**). The lowered standard deviations confirmed that both complexes reached convergence in time period. And therefore, for more precise comparison, the calculations of MM/PBSA binding energy only took account of simulation snapshots from the 125^th^ ns to 200^th^ ns (**Figure 8B**). In conjunction with the C-alpha backbone RMSD of the complexes but contrary to the hypothesis, FGF5-H174 possessed stronger binding affinity (of 28.8 kJ/mol) than that of FGF5-WT, which was 130.4 kJ/mol (**Figure 8B**). The results of these findings prompt the question: could the reduced activity of FGF5-H174 at FGFR1 previously observed be attributed to altered residual connectivity or binding mode of the complexes? Therefore, the free energy landscapes were used to extract and compare meta-stable states (**Figure 9**), which were subsequently analyzed with interaction networks (**Figure 10**) for better insights into this phenomenon. The free energy landscapes demonstrated that the conformational movement of FGFR1-FGF5-WT complex was more scattered and predicted that more Gibbs free energy would be required for this complex to reach its meta-stable state from high-energy state (of 8.17 kJ/mol) (**Figure 9A, top panel**) compared to that of FGFR1 - FGF5-H174 complex, which only required 7.54 kJ/mol (**Figure 9B, top panel**). Meta-stable states were further extracted from the free energy landscapes and structurally compared through structural alignment (**Figure 9A and B, bottom panels**). It was found that the FGFR1 proteins had altered conformations of 3 loop segments, which were N296 to L306, L328 to D336, and V267 to V273 (**Figure 9A and B, bottom panels**). But most importantly, FGF5-WT managed to “flip” the segment V267 to V273 outward, whereas this segment remained inward for FGF5-H174 (**Figure 9A and B, bottom panels**). This segment is known to hold important functions, such as acting as an anchor for the KL2 domain of the α-klotho to induce phosphate homeostasis and vitamin D metabolism (Buchanan et al., 2020; G. Chen et al., 2018). Therefore, it is also reasonable to suspect if the residue connectivity or binding more could be altered due to the missense mutation, and residue interaction networks of the meta-stable states were constructed to further investigate this suspicion (**Figure 10A and B**). Overall, the residue interaction networks of FGFR1 - FGF5-WT and - FGF5-H174 complexes had 480 and 430 edges/interactions, respectively (**Figure 10A, B, and C**). Particularly, almost all of interaction types decreased following the missense mutation, except inter-chain pi-pi stack and ionic inter-chain interactions (**Figure 10C**). Additionally, to investigate if there were any alteration in hotspot residues, the distribution histograms of the degrees of nodes (or residues) were constructed and demonstrated that FGFR1 - FGF5-WT and - FGF5-H174 complexes had a peak at degree of 1 and 2, respectively (**Figure 10D and E**). However, only FGFR1 - FGF5-WT complex had a distinctive hump at degree of 8, an indication that the missense mutation reduced the amount of hotspot residues as well as potentially altering the location of these residues. Residues with 8 or more degrees are listed in **Figure 10F**, confirming that not only the amount of hotspot residues had been reduced but also that the hotspot locations had been switched. For example, FGFR1-F197 with the largest amount of degrees had been replaced with F176; FGF5-F141 with 9 degrees had been replaced with H214, or only FGFR1-F197 and FGF5-F124 remained in both columns (**Figure 10F**).

**Figure 8.**
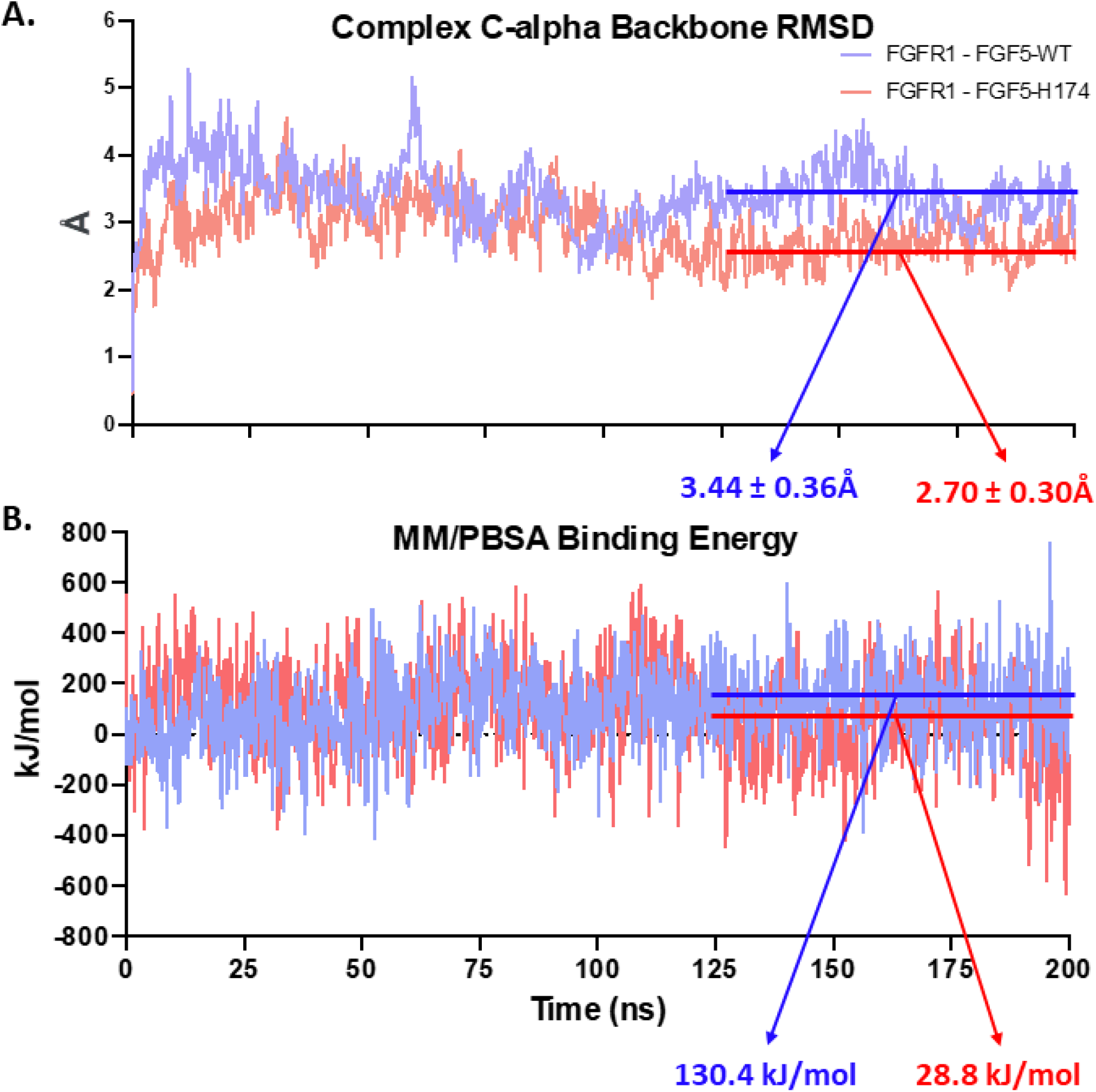
Molecular Dynamics Simulations of FGF5-WT and FGF5-H174 in complex with FGFR1. (A) C-alpha backbone RMSD of the complexes. (B) MM/PBSA binding energies of FGF5 proteins towards FGFR1.

**Figure 9.**
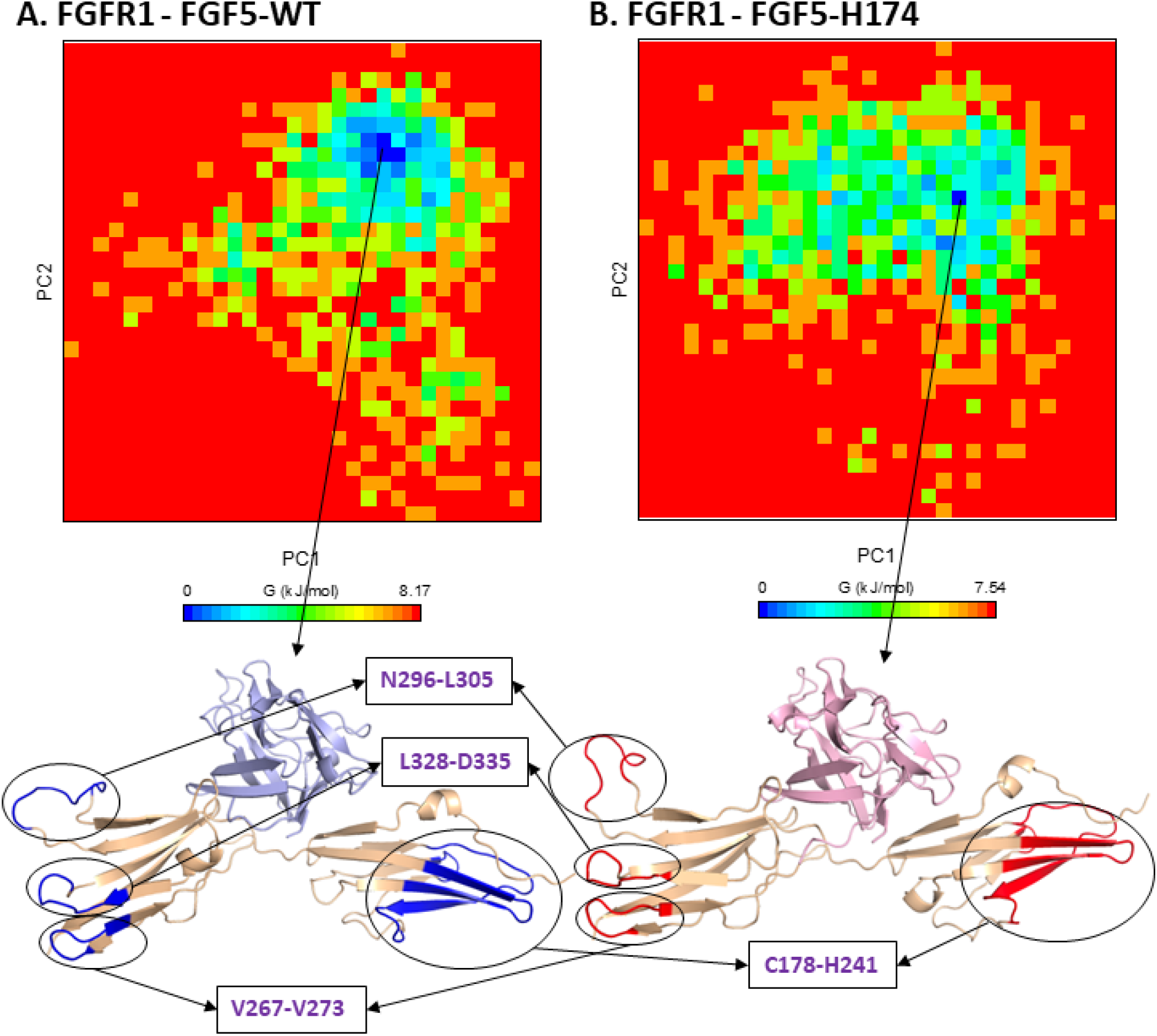
Free energy landscapes of FGFR1-FGF5-WT and - FGF5-H174 in complexes meta-stable states derived from the C-alpha backbone principal component analysis (PCA). (A) FGFR1 - FGF5-WT. (B) FGFR1 - FGF5-H174.

**Figure 10.**
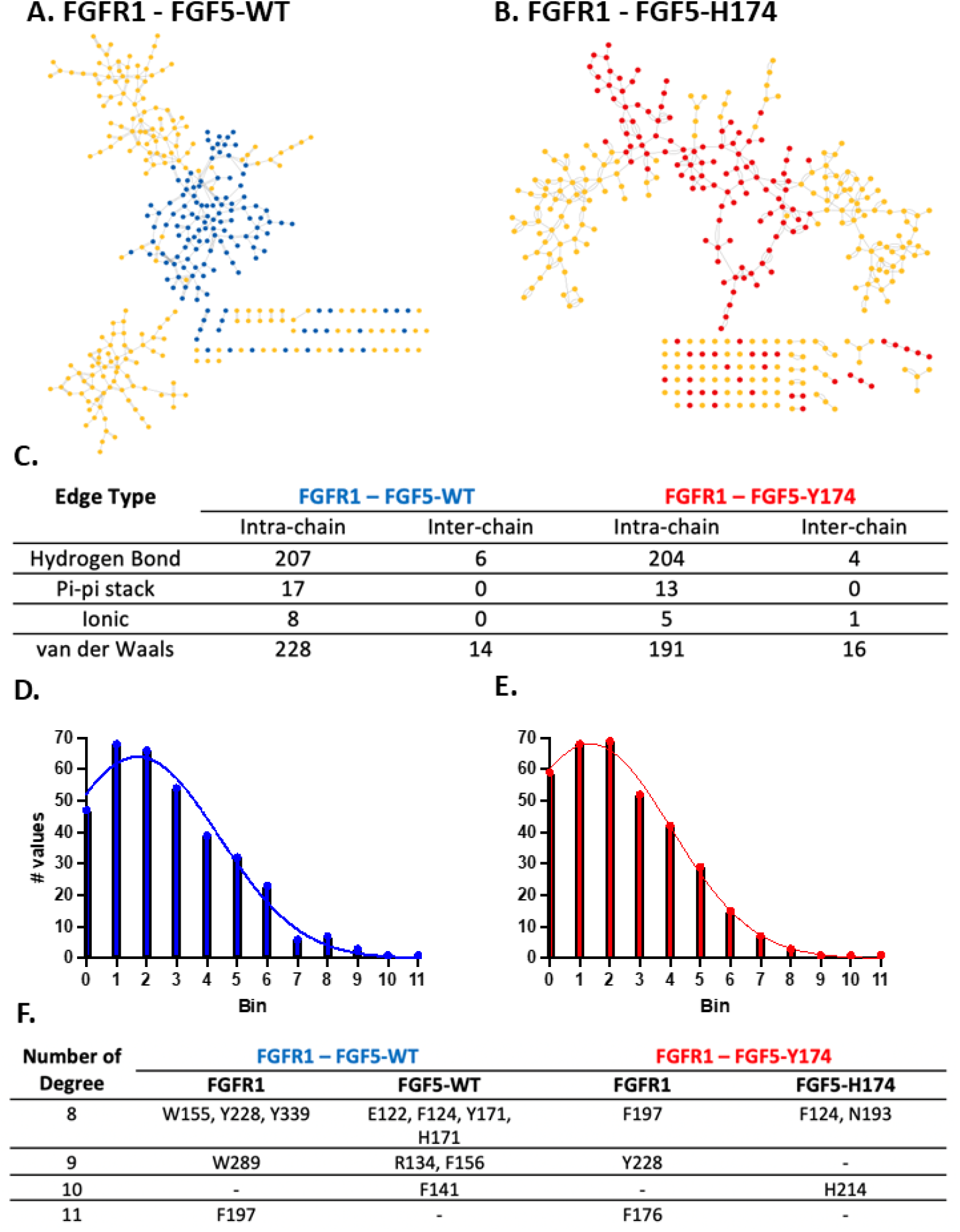
Residue interaction network analysis. Networks of residue interactions for (A) FGF5 - WT-FGFR1 and (B) FGF5 - Y174H-FGFR1. (C) Panel summarizing the total intra-chain and inter-chain interactions. Histograms depicting changes in the distributions of degree for (D) FGF5-WT-FGFR1 and (E) FGF5-Y174H-FGFR1. (F) Panel summarizing specific residues with degrees of 8 or larger.

## Discussion

The analysis of the molecular dynamics simulations confirmed the hypothesis that the missense mutation Y174H of FGF5 will contribute to changes in protein behaviors and structures. Particularly, interactions with the solvent, secondary structures, conformational stability, as well as connectivity with its primary receptor FGFR1 were evaluated. The mutation resulted in an increase in SASA (a biophysical marker for pathogenic protein (Savojardo et al., 2020)), further reinforced a decrease in number of hydrogen bonds within the protein and an increase in number of hydrogen bonds between the protein and solvent. Furthermore, the mutation saw a decrease in sheet but an increase in coil structures. Specifically, secondary changes were highly observable at residues 146 to 152 as well as 160 to 171. Interestingly, these residues are also in proximity with residues 91, 123, 131, 133, 162, 164, 172, 190, and 215 where H174 showed decreased and sporadic interactions compared to Y174, confirming the importance of this residue, conserved throughout multiple species (Higgins et al., 2014). Besides, analyses of the C-alpha backbone RMSD, the number of salt-bridges, and protein residue RMSF demonstrated that: compared to its wild-type counterpart, FGF5-H174 experienced more movement, conformational changes, and fluctuations – accompanied by a decrease in the number salt-bridges, an indicator of structural stability (Bosshard et al., 2004). Lastly, we employed protein-protein docking to explore the effect of this mutation on FGFR1 binding, which were further examined with molecular dynamics simulations and residue interaction network.

Contrary to the initial prediction that the missense mutation would lead to unstable binding, MM/PBSA binding energies demonstrated that FGF5-H174 had a stronger binding affinity towards FGFR1. However, it could be speculated that FGF5-Y174H previously demonstrated a decreased inhibition on FGFR1 (Higgins et al., 2014) because of its difference in residue signaling and connectivity. The residue interaction network analysis on the meta-stable states of the complexes confirmed this speculation. To the best of my knowledge, this is the first study to directly investigate the structure and dynamics of FGF5 as well as its missense mutation variant FGF5-Y174H, underlying trichomegaly. It is recommended that further *in vitro* and biophysical methods to validate these findings.

## References

Allerstorfer, S., Sonvilla, G., Fischer, H., Spiegl-Kreinecker, S., Gauglhofer, C., Setinek, U., Czech, T., Marosi, C., Buchroithner, J., Pichler, J., Silye, R., Mohr, T., Holzmann, K., Grasl-Kraupp, B., Marian, B., Grusch, M., Fischer, J., Micksche, M., & Berger, W. (2008). FGF5 as an oncogenic factor in human glioblastoma multiforme: Autocrine and paracrine activities. Oncogene, 27(30), 4180–4190. https://doi.org/10.1038/onc.2008.61

Bosshard, H. R., Marti, D. N., & Jelesarov, I. (2004). Protein stabilization by salt bridges: Concepts, experimental approaches and clarification of some misunderstandings. Journal of Molecular Recognition: JMR, 17(1), 1–16. https://doi.org/10.1002/jmr.657

Buchanan, S., Combet, E., Stenvinkel, P., & Shiels, P. G. (2020). Klotho, Aging, and the Failing Kidney. Frontiers in Endocrinology, 11, 560. https://doi.org/10.3389/fendo.2020.00560

Chen, B., Hu, R., Min, Q., Li, Y., Parkinson, D. B., & Dun, X.-P. (2020). FGF5 Regulates Schwann Cell Migration and Adhesion. Frontiers in Cellular Neuroscience, 14, 237. https://doi.org/10.3389/fncel.2020.00237

Chen, G., Liu, Y., Goetz, R., Fu, L., Jayaraman, S., Hu, M.-C., Moe, O. W., Liang, G., Li, X., & Mohammadi, M. (2018). α-Klotho is a non-enzymatic molecular scaffold for FGF23 hormone signalling. Nature, 553(7689), 461–466. https://doi.org/10.1038/nature25451

Clementel, D., Del Conte, A., Monzon, A. M., Camagni, G. F., Minervini, G., Piovesan, D., & Tosatto, S. C. E. (2022). RING 3.0: Fast generation of probabilistic residue interaction networks from structural ensembles. Nucleic Acids Research, 50(W1), W651–W656. https://doi.org/10.1093/nar/gkac365

Galindo-Murillo, R., Robertson, J. C., Zgarbová, M., Šponer, J., Otyepka, M., Jurečka, P., & Cheatham, T. E. (2016). Assessing the Current State of Amber Force Field Modifications for DNA. Journal of Chemical Theory and Computation, 12(8), 4114–4127. https://doi.org/10.1021/acs.jctc.6b00186

Han, D., Wang, M., Yu, Z., Yin, L., Liu, C., Wang, J., Liu, Y., Jiang, S., Ren, Z., & Yin, J. (2019). FGF5 promotes osteosarcoma cells proliferation via activating MAPK signaling pathway. Cancer Management and Research, 11, 6457–6466. https://doi.org/10.2147/CMAR.S200234

Higgins, C. A., Petukhova, L., Harel, S., Ho, Y. Y., Drill, E., Shapiro, L., Wajid, M., & Christiano, A. M. (2014). FGF5 is a crucial regulator of hair length in humans. Proceedings of the National Academy of Sciences of the United States of America, 111(29), 10648–10653. https://doi.org/10.1073/pnas.1402862111

Iwabu, J., Yamashita, S., Takeshima, H., Kishino, T., Takahashi, T., Oda, I., Koyanagi, K., Igaki, H., Tachimori, Y., Daiko, H., Nakazato, H., Nishiyama, K., Lee, Y.-C., Hanazaki, K., & Ushijima, T. (2019). FGF5 methylation is a sensitivity marker of esophageal squamous cell carcinoma to definitive chemoradiotherapy. Scientific Reports, 9(1), 13347. https://doi.org/10.1038/s41598-019-50005-6

Land, H., & Humble, M. S. (2018). YASARA: a tool to obtain structural guidance in biocatalytic investigations. In Protein Engineering (pp. 43–67). Springer.

Li, W.-R., Liu, C.-X., Zhang, X.-M., Chen, L., Peng, X.-R., He, S.-G., Lin, J.-P., Han, B., Wang, L.-Q., Huang, J.-C., & Liu, M.-J. (2017). CRISPR/Cas9-mediated loss of FGF5 function increases wool staple length in sheep. The FEBS Journal, 284(17), 2764–2773. https://doi.org/10.1111/febs.14144

Ornitz, D. M., & Itoh, N. (2015). The Fibroblast Growth Factor signaling pathway. Wiley Interdisciplinary Reviews. Developmental Biology, 4(3), 215–266. https://doi.org/10.1002/wdev.176

Punzalan, F. E. R., Cutiongco-de la Paz, E. M. C., Nevado, J. J. B., Magno, J. D. A., Ona, D. I. D., Aman, A. Y. C. L., Tiongson, M. D. A., Llanes, E. J. B., Reganit, P. F. M., Tiongco, R. H. P., Santos, L. E. G., Aherrera, J. A. M., Abrahan, L. L., Agustin, C. F., Bejarin, A. J. P., & Sy, R. G. (2022). The rs1458038 variant near FGF5 is associated with poor response to calcium channel blockers among Filipinos. Medicine, 101(5), e28703. https://doi.org/10.1097/MD.0000000000028703

Ren, Y., Jiao, X., & Zhang, L. (2018). Expression level of fibroblast growth factor 5 (FGF5) in the peripheral blood of primary hypertension and its clinical significance. Saudi Journal of Biological Sciences, 25(3), 469–473. https://doi.org/10.1016/j.sjbs.2017.11.043

Rossetto, J. D., Nascimento, H., Muccioli, C., & Belfort, R. (2013). Essential trichomegaly: Case report. Arquivos Brasileiros De Oftalmologia, 76(1), 50–51. https://doi.org/10.1590/s0004-27492013000100015

Savojardo, C., Manfredi, M., Martelli, P. L., & Casadio, R. (2020). Solvent Accessibility of Residues Undergoing Pathogenic Variations in Humans: From Protein Structures to Protein Sequences. Frontiers in Molecular Biosciences, 7, 626363. https://doi.org/10.3389/fmolb.2020.626363

Xin, Q., Han, Y., Jiang, W., Wang, J., Luan, Y., Ji, Q., & Sun, W. (2022). Genetic susceptibility analysis of FGF5 polymorphism to preeclampsia in Chinese Han population. Molecular Genetics and Genomics: MGG, 297(3), 791–800. https://doi.org/10.1007/s00438-022-01889-z

Yan, Y., Zhang, D., Zhou, P., Li, B., & Huang, S.-Y. (2017). HDOCK: A web server for proteinprotein and protein-DNA/RNA docking based on a hybrid strategy. Nucleic Acids Research, 45(W1), W365–W373. https://doi.org/10.1093/nar/gkx407

